# Sperm of more colorful males are better adapted to ovarian fluids in lake char (Salmonidae)

**DOI:** 10.1101/2021.10.02.462848

**Authors:** David Nusbaumer, Laura Garaud, Christian de Guttry, Laurie Ançay, Claus Wedekind

## Abstract

Fish often spawn eggs with ovarian fluids that have been hypothesized to support sperm of some males over others (cryptic female choice). Alternatively, sperm reactions to ovarian fluids could reveal male strategies linked to their likely roles during spawning. Sperm of males who would usually be close to females during spawning are then expected to be better adapted to the presence of ovarian fluids than to water only, while the reverse would be expected for males that typically spawn at larger distance to the females. We tested these predictions with gametes and ovarian fluids from wild-caught lake char (*Salvelinus umbla*). We found that sperm of more colorful males showed increased sperm velocity in diluted ovarian fluids while sperm of paler males were fastest in water only. We then let equal numbers of sperm compete for fertilizations in the presence or absence of ovarian fluids and used microsatellite markers to assign in total 1,464 embryos (from 70 experimental trials) to their fathers. Overall, sperm of more colorful males reached higher fertilization success than sperm of pale males. This difference was enhanced by the presence of ovarian fluids and best explained by the increased sperm velocity. Sperm competitiveness was not enhanced with decreasing genetic distance to a given female, although parallel stress tests on embryos had revealed that females would profit more from mating with least related males rather than most colored ones. We conclude that sperm of more colorful males are best adapted to ovarian fluids, and that the observed reaction norms reveal male strategies rather than cryptic female choice.

## Introduction

Ejaculate economics predicts that males invest strategically into sperm according to the expected microecology they will experience after ejaculation (Parker & Pizzari 2010). This microecology is strongly defined by competitors and the level of sperm competition. Males are predicted to increase sperm number and/or sperm velocity with increasing risk of sperm competition (Parker & Pizzari 2010). This prediction is meanwhile well supported in many taxa (Magris 2021). Another aspect of the microecology that sperm have to deal with is the biochemical environment that females create around their eggs with their reproductive fluids. The functional significance of such female reproductive fluids is not well understood yet (Gasparini *et al*. 2020). In externally spawning fishes, the fluids with which females expel their eggs are called “ovarian fluids” and consist of various inorganic and organic compounds (Lahnsteiner *et al*. 1995).

Salmonids are externally spawning fish whose ovarian fluids vary in amount and composition among species and even within populations (Lahnsteiner *et al*. 1995; Zadmajid *et al*. 2019). Viscosity of undiluted ovarian fluids is about 2 to 3 times higher than that of water (Zadmajid *et al*. 2019), but ovarian fluids will quickly be diluted when expelled with eggs into water. During spawning, freshly spawned eggs and ovarian fluids are swirled around by male and female movements so that some eggs may no more be surrounded by significant amounts of ovarian fluids at the time of fertilizations while others still are. A likely predictor of how much sperm will be exposed to ovarian fluids is the position of the male during spawning. Arguably the closer a male is with its urogenital opening to a female’s vent during egg release, the more likely its sperm will be exposed to ovarian fluids.

Ovarian fluids have repeatedly been found to support sperm velocity, motility, and longevity (Zadmajid *et al*. 2019), i.e. key targets of sexual selection (Evans *et al*. 2013). They have also been discussed as allowing for certain forms of cryptic female choice (Firman *et al*. 2017). Their release with eggs can, for example, specifically promote paternity of dominant males over parasitic spawners (“sneaker males”) (Alonzo *et al*. 2016; Egeland *et al*. 2016; Makiguchi *et al*. 2016; Lehnert *et al*. 2017). The support of sperm function can be even more specific: Ovarian fluids of guppies (*Poecilia reticulata*) were found to slow down sperm of full-sibs as compared to unrelated males, potentially to reduce inbreeding (Gasparini & Pilastro 2011). The presence of ovarian fluids during spawning has also been found to reduce inbreeding and hence offspring mortality by differentially supporting some sperm over others in Chinook salmon (*Oncorhynchus tshawytscha*) (Rosengrave *et al*. 2016; Gessner *et al*. 2017; Lehnert *et al*. 2017), while ovarian fluids of lake trout (*Salvelinus namaycush*) were found to promote sperm velocity of more related males (Butts *et al*. 2012), potentially to reduce outbreeding depression in this salmonid. Indeed, Yeates et al. (2013) found that ovarian fluids specifically support sperm of the same *Salmo* species over sperm of another *Salmo* species, i.e. they promote reproductive isolation by a species-specific chemoattraction. All these examples have mainly been discussed in the light of adaptive female reproductive strategies. However, many of the observed effects of ovarian fluids on sperm function could reveal variation in male reproductive strategies, as predicted by the theory of ejaculate economics and experimentally demonstrated in other vertebrates (Jeannerat *et al.*).

Salmonid males can play various roles during spawning (Esteve 2005), and many of these roles have been found to affect milt and sperm characteristics (Magris 2021). In migratory populations, for example, there is usually a fraction of males that avoid the costs of migration but then remain small and subdominant to their migratory rivals at spawning. They therefore reproduce mainly surreptitiously and invest disproportionally into sperm number and sperm velocity in order to increase the competitiveness of their sperm (Young *et al*. 2013). Evidence for strategic male investments has also been found within non-migratory populations. Male dominance at the spawning place is usually size dependent (Esteve 2005), and older males and larger male whitefish (*Coregonus zugensis*) have indeed been found to invest less into gonad weight and to have slower sperm than young and small males who typically spawn further away from the female’s vent (Rudolfsen *et al*. 2008). However, the links between male size and sperm traits can vary among populations (Perroud *et al*. 2021) and be plastic in response to the perceived social environment (Bartlett *et al*. 2017). When Rudolfsen et al. (2006), for example, controlled for size effects by confining pairs of size-matched male Arctic char (*Salvelinus alpinus*) in an enclosure and let them develop a dominance hierarchy, sperm characteristics changed quickly: as predicted from ejaculate theory, dominance led to reduced sperm velocity. Egeland et al. (2016) then used an analogous setup to show that sperm of dominant males are better designed to swim in ovarian fluids. One possible explanation for this increased performance is that ovarian fluids somehow recognize sperm of dominant males and promote them specifically (cryptic female choice). Alternatively, dominant males can expect to be close to the females during gamete release and may therefore produce sperm that do well when exposed to ovarian fluids. If so, any other male trait that is likely to affect the distance between male and female during spawning would be predicted to affect sperm reaction to ovarian fluids. Here we use wild-caught lake char (*Salvelinus umbla*) from Lake Geneva (Switzerland) to test whether male sexual ornamentation is such a trait.

The lake char is a non-migratory salmonid endemic to Alpine lakes. It spawns in winter in lek-like mating system where male competition is usually intense. This species is closely related to the Arctic char (Kottelat & Freyhof 2007) that has often been used to study effects of ovarian fluid on sperm motility (e.g.Turner & Montgomerie 2002; Urbach *et al*. 2005; Egeland *et al*. 2016). Both species develop spawning colorations. Arctic char often develop strong red colorations in both sexes (Janhunen *et al*. 2011). Male lake char of our study population are mostly yellow during spawning season while females hardly develop any spawning colorations.

In other fish families, sexual ornaments are often positively linked to aspects of sperm performance (Kortet *et al*. 2004; Mehlis *et al*. 2013; Fukuda & Karino 2014; Kekäläinen *et al*. 2014), probably because both types of traits can react to induced stress (Kekäläinen *et al*. 2014), suggesting that high quality sperm in well ornamented males is an indicator of health and vigor rather than of alternative mating strategies. The pattern in salmonids seems less clear. Janhunen et al. (2009) found in a captive population of Arctic char that the intensity of male coloration was positively correlated with sperm velocity (when tested in water) while dominant males are expected to have reduced sperm velocity (Rudolfsen *et al*. 2006). Combined, these findings suggest that male dominance and coloration are not necessarily linked in char. Nevertheless, if females prefer colorful males, we predict that not only dominant but also colorful males are usually closer to females during spawning than other types of males. If so, sperm of colorful males are predicted to do better when exposed to ovarian fluids than sperm of less colorful males.

Here we test whether (i) male coloration of lake char predicts sperm performance as it does in other fish, (ii) sperm of colorful males react better to presence of ovarian fluids that sperm of less colorful males, as predicted if colorful males are preferred and/or more dominant and therefore able to spawn closer to females, (iii) colorful males reach disproportional high fertilization success in sperm competition, (iv) the presence of ovarian fluids change the outcome of sperm competition because of its effects in sperm performance, and (v) ovarian fluids of different females differ in their effects on sperm. We also test whether (vi) coloration is an indicator of male inbreeding coefficient, and if not, whether (vii) male inbreeding coefficient or (viii) their genetic relatedness to females affect sperm characteristics and the outcome of sperm competition in water or in diluted ovarian fluids.

## Methods

### Sampling and handling of fish, gametes, and ovarian fluids

Wild char were caught in December 2017 and 2018 from three spawning sites in Lake Geneva (Ripaille, 46.3957N-6.4718E; Locum, 46.4049N-6.7588E; Meillerie, 46.4113N-6.7216E) by local fishermen using gill nets (48 mm mesh size, set over night) and transported to a research facility (INRA, Thonon-les-Bains, France) were they were kept in a 1,000 L circular tank with free-flowing lake water before further processing. In 2017, 10 males and 4 females were haphazardly chosen for the experiment. Males were processed on the first day after catching and females on the second day. Each fish was anaesthetized in 10 L water containing 3 mL of Eugenol (clove oil in ethanol at a 1:9 ratio), photographed in a custom-made photobox under standardized conditions (17 mm, f/5.0, 1/200 s, ISO 400, WB 4000 K, JPG 24 Mpx), and striped to collect gametes. Tissue samples were taken from the anal fin and stored in 70% ethanol at −20°C for further analysis. In 2018, further 24 males were sampled and handled using the same procedures.

Milt was directly striped into 50 mL conical tubes (Falcon, BD Biosciences, Allschwil, Switzerland), carefully avoiding contamination by urine, feces, or water. Between 2 and 5 mL of milt were obtained per male. These samples were stored at 4°C (< 2 hours) until further use. When all males had been stripped, 1.5 mL of each milt sample was diluted 1:9 in Storfish (IMV Technologies, France; an inactivating medium) in 15 mL tubes (Falcon, BD Biosciences, Allschwil, Switzerland). These tubes were then placed on ice and are referred to as ‘diluted milt’.

Eggs and ovarian fluids were stripped into individual plastic containers. About 20 mL of ovarian fluid each was separated from the eggs with a syringe and stored in 50 mL tubes at 4°C. Eggs were kept (<1h) in the dark at 4°C in about 20 mL Ovafish (IMV Technologies, France) to prevent drying.

### Color measurement

Custom-made macros in Fiji were used to analyze male skin colorations. First, the white balance of each image was standardized with the help of the white and black values of the color scale. Skin gray value was then determined as the overall body gray value in the RGB color space (range from 0 = black, to 255 = white; using mean of all pixels in each of the RGB channels). Fish redness and yellowness were measured from 3 squares (about 1% of the total body area each) in the pectoral region, the ventral region and the anal region, as described by Parolini et al. (2018) (Fig. S1). Redness and yellowness were measured as the *a\** and *b\** components of the CIE-*L*a*b\** color space, respectively (the a* axis for greenness-redness and the b* axis for blueness-yellowness both range from −100 to 100). All measures of yellowness were significantly correlated (Pearson’s *r* > 0.92, *p* always < 0.001). Similarly, all measures of redness were significantly correlated (Pearson’s *r* > 0.75, *p* < 0.013). Therefore, the means of the three measurements per fish were used for each color. Gray values were correlated with yellowness (the yellower the lighter) and with redness (the redder the darker; Fig. S1) and were therefore ignored in further analyses.

### Sperm characteristics of freshly caught fish

Sperm characteristics were measured on the day of sampling with Qualisperm® (AKYmed, Switzerland). Briefly, 20 μL of the diluted milt was activated in a microtube with 980 μL ice-cooled standardized water (OECD 1992) and vortexed for 5s. Then, 2 μL of this activated solution was pipetted into the well of a 4-well chamber slide (Leja, AKYmed, Switzerland), on a cooling station set at 4.5°C (the expected water temperature at the spawning ground). Sperm concentration, sperm velocity (average path velocity), and percentage of immotile sperm (sperm motility) were recorded under a phase contrast microscope at 20X magnification 20 s after activation in 2017 and 25 s after activation in 2018. Sperm longevity was recorded with a stopwatch from the time of activation until no progressive sperm motion could be observed in the frame of capture. Two replicated measures were taken for each sperm sample to test for consistency, and means of these measurements were used for statistics. The concentration of active spermatozoa was calculated for each male by multiplying the total sperm concentration with the motility (e.g. 2,000 Mio/mL times 80% motility = 1,600 Mio/mL active spermatozoa).

### Hematological analyses

Hematological analyses are frequently used to determine stress in fish (Seibel *et al*. 2021). High leukocyte counts and especially also relative lymphocytosis (percentage of lymphocytes among leukocytes) have been associated with acute infections, e.g. in eel (*Anguilla anguilla*) infected with the virulent nematode *Anguillicola crassus* (Haenen *et al*. 2010), or have been found in response to exposure to pesticides (Oluah *et al*. 2020), and even in anticipation of predation risk (Meuthen *et al*. 2020). Thrombocytosis (elevated platelet counts) is another possible indicator of an acute infection (Rose *et al*. 2012; Yan *et al*. 2013).

In order to determine these potential stress indicators, a blood sample (2-3 mL) was drawn from the caudal vasculature of males with a 3 mL syringe mounted with a 23 G needle heparinized with heparin 1 IU/μL. The blood was emptied in 2 mL microtubes, centrifuged 10 min at 10,000 g and 4°C. The plasma was then carefully collected, disposed in 1.5 mL microtubes and frozen in the vapor of liquid nitrogen for transportation and stored at −80°C in the lab for further analysis. A drop a blood from the syringe was used to make two blood smears per individual, on microscope frosted slides. Blood smears were dried in the field and fixed for 3 minutes in absolute ethanol back in the lab. Slides were stained for 20 minutes in Giemsa stain and rinsed twice in PBS. From each slide, 10 photographs (i.e. 20 per individual) were taken on an EVOS XL Core inverted light transmission microscope (Thermofisher, Switzerland) at 40x magnification. Counts of thrombocytes, lymphocytes, and granulocytes were done manually by a naïve observer (Fig. S2). Counts of total number of cells were done automatically with a custom-made macro in Fiji (Fig. S2).

### Experiment 1: Sperm characteristics in response to ovarian fluids

For the 2017 sample, the same protocol was used on the day of experimental fertilizations, i.e. one day after catching the fish from the wild, to determine each male’s sperm characteristics in each female’s ovarian fluid (40 combinations of milt samples and ovarian fluids) and in water (controls, 4 times replicated each; Table S1). To do so, 10 μL of the diluted milt were activated either with 490 μL standardized water or with 490 μL ovarian fluid solution. The concentration of ovarian fluid in this solution was 50% as this would reflect a possible concentration of ovarian fluid in the water when sperm swim towards the egg (Rosengrave *et al*. 2016).

### Experiment 2: Competitive fertilization trials and monitoring of embryos

The 10 males were haphazardly assigned to 5 dyads (pairs of competitors) whose sperm then competed for fertilization of 24 eggs/female, either activated in water only or in water with ovarian fluid. Each possible combination of dyad x female was tested in both environments twice, resulting in 80 sperm competition trials (5 dyads x 4 females x 2 treatments x 2 replicates; Table S2) that involved in total 1,920 eggs. However, due to an accident during handling, 10 of these 80 experimental cells were lost (all of the same female, prepared to be exposed to sperm activated in water only, see Table S2), reducing the number of eggs that could be monitored to N = 1,680.

In preparation of these sperm competition trials, the eggs of each female were first washed twice with 200 mL Ovafish® (IMV Technologies, France) to remove ovarian fluids. Then, 20 batches of 24 eggs each were placed in wells of 6-well plates (Falcon, BD Biosciences, Allschwil, Switzerland). Ten mL of diluted milt of 2 males each was prepared such that each male was represented with the same concentration of active sperm (25 Mio/mL) in the mix. One mL of each mix was then used to fertilize the 2 batches of 24 eggs/female. This 1 mL of mix was activated in a separated tube with either 4 mL of standardized water or 4 mL of standardized water with ovarian fluid (ratio ovarian fluid to water = 1:2) and vortexed for 5s. The solution was poured in a well with eggs. Two minutes later, i.e. after fertilization could be expected to have happened, standardized water was added to fill the wells (16.8 mL) and the eggs were left undisturbed in a dark environment for 2 hours to allow hardening.

Each batch of 24 supposedly fertilized eggs was transferred to a 50 mL tube filled with standardized water and transported on ice to a climate chamber where they were rinsed for 30s under running tap water (4 L/min) in a sterilized tea strainer. The eggs were then distributed singly to wells of 24-well plates (Falcon, BD Biosciences, Allschwil, Switzerland) filled with 1.8 mL of autoclaved standardized water (OECD 1992). Embryos were raised at 4.5°C in a 12h:12h light-dark cycle. Fertilization success was assessed at the neurula stage (14 days post fertilization). No visible embryo at this stage was interpreted as non-fertilized egg. Embryo mortality, the timing of hatching, hatchling size, and hatchling growth was monitored from then on in the course of two parallel studies, one that combines these data with further stress experiments (by a pathogen and a chemical pollutant) on embryos produced in a regular full-factorial breeding design to test for parental effects on embryo performance (Garaud *et al*. 2022), and another one on sex-specific stress tolerance (Nusbaumer *et al*. 2021). Embryos and larvae that died during the observational period were transferred to individual microtubes and stored in 96% ethanol at −20°C for further analysis. Larvae were euthanized 14 days after hatching with a spike of 100 μL of Koi Med® Sleep 4.85% (Koi & Bonsai Zimmermann, Germany) in the well before being transferred to individual microtubes.

### Genotyping and paternity assignment based on microsatellite markers

In the case of larvae and adults, DNA was extracted from up to 25 mg tissue with an extraction robot and the DNAeasy Blood & Tissue kit (Qiagen, Hilden, Germany) following manufacturer’s instructions. Unhatched embryos were homogenized and DNA was extracted using the Qiamp Fast DNA Stool mini kit protocol (Qiagen, Germany) as in Wilkins et al. (2015). A first multiplex PCR of 3 polymorphic microsatellite markers (Savary *et al*. 2017) could be used to assign most embryos and larvae to their fathers (Bernatchez & Duchesne 2000). A second multiplex PCR of 3 further microsatellite markers (Savary *et al*. 2017) was used for to assign the remaining offspring (Table S3). In the first multiplex PCR, the *sdY* sex marker (Yano *et al*. 2013) was included to sex offspring (Table S3) for a parallel study on sex-specific stress tolerance (Nusbaumer *et al*. 2021). Multiplex PCR were run on a Biometra thermocycler in 20 μL reaction volume using HotStarTaq DNA Polymerase (Qiagen, Germany) reagents (i.e. 2 μL PCR buffer 10X, 4 μL QSolution 5X, 0.4 μL HotStar Taq DNA Polymerase), 0.4 μL of dNTPs 10 mM and 0.4 μL of each primer). PCR cycles were run as follow: initial heat activation of 15 min at 95 °C; 35 cycles of 30 s denaturation at 94°C, 90 s annealing at 57°C and 60 s extension at 72°C; followed by 30 min of final extension at 72°C and 10 min at 4°C. DNA extracted from unhatched embryo was amplified using the same protocol, but in 10 μL reaction volume and with 40 cycles. PCR products were diluted 2X and run on an ABI3100 sequencer (Applied Biosystems, USA). Genotypes were read using Genemapper v4.0 (Applied Biosystems, USA). Genotyping of 3 dead embryos was not successful possibly due to poor DNA quality, bringing the final number of genotyped offspring to N = 1,475.

Adults were genotyped twice (no discrepancies observed). Paternity was assigned using CERVUS v3.0 (Kalinowski *et al*. 2007) as recommended for categorical assignments in controlled experiments (Jones *et al*. 2010). Input simulation parameters were set with 6 loci, 10,000 offspring, 2 candidate fathers with a proportion of sampling of 1 and a proportion of typing of 1, a minimum of 2 typed loci and an allelic dropout/mistyping rate of 0.05 as recommended by Wang (2004), and an assignment confidence of 95%.

### Double-digested RAD sequencing and SNPs calling

DNA from all males and females of experiment 2 was used to also determine male multi-locus heterozygosity (MLH) and the genomic similarity (“kinship”) between males and females of all possible combinations (N = 40). DNA concentration was measured using Qubit 2.0 (Thermo Fisher Scientific, Waltham, Massachusetts) while its integrity was verified on a 1% agarose gel. DNA was subsequently standardized to 20 ng/μl. Two sequencing libraries were produced using all 10 males and 4 females each following the Brelsford et al. (2016) protocol adapted from Parchman et al. (2012). Briefly, DNA was digested using the enzymes EcoRI-HF and MspI (New England Biolab, Ipswich, Massachusetts, United States) and a unique EcoRI barcode was ligated to each individual. After library purification, PCR amplification were performed and fragments size-selected in between 400-550bp. Libraries were then single-end genotyped on 2 lanes of Illumina HiSeq 2500 with fragments of 125 bp length at Lausanne Genomic Technologies Facility (University of Lausanne).

After the quality control on the resulting fastq files done with FASTQC v0.11.7 (Andrews 2010), reads were trimmed to 110bp given the low per-base quality of the sequences (<20 Phred score). The resulting RAD data were analysed using Stacks 1.48 (Catchen *et al*. 2013) using the default parameters unless otherwise specified. Individuals sequences were demultiplexed using *process_radtags* (Stacks 1.48) and reads mapped to the *Salvelinus spp* reference genome (Christensen *et al*. 2018) using BWA (Li & Durbin 2010). *Pstacks* (Stacks 1.48) was done selecting a minimum stack depth of 5 (*-m* 5) and using the bounded SNP model. The catalogue of loci was created using *Cstacks* (Stacks 1.48) allowing 2 mismatches (*-n* 2) between loci. *Populations* (Stacks 1.48) was used to generate the VCF file considering only loci present in 100% of the individuals and markers heterozygosity of 0.5 (*--max_obs_het* 0.50). None of the individuals was excluded based on genotyping rate. Further filtering was done using *vcftools*, v0.1.15 (Danecek *et al*. 2011). To reduce incorrect heterozygosity call and remove paralogs, loci were filtered for a minimum coverage of 10X (*--min-meanDP* 10) and a maximum coverage of 50X (*--max-meanDP* 50; two times the mean coverage). Only loci under Hardy-Weinberg equilibrium were retained with a P-value treshold of 0.05 (*--hwe* 0.05). Finally, all the loci that were not shared between all individuals were discarded (*--max-missing* 1). No filtering was made based on minor allele frequencies. A total of 4,150 SNPs was retained after filtering with a mean coverage per individual of 29X.

Individual MLH, i.e. the number of heterozygous loci on the genotyped portion of the genome, was calculated using *vcftools (--het*) (Danecek *et al*. 2011). The kinship, i.e. the expected inbreeding coefficient for the progeny of one family, was estimated using the *beta.dosage* function of the package *Hierfstat* (Goudet 2005). This function generates a genomic matrix based on allele dosage to estimate kinship coefficients considering the proportion of alleles shared between individuals (Goudet *et al*. 2018). The inbreeding coefficients for females and males, *F_beta_*, were obtained by extracting the diagonal of the genomic matrix obtained with *beta.dosage*. Fig. S3 shows how MLH (range in males: 0.242 to 0.259) relates to *F_beta_* (range in males: 0.029 to 0.092). Further analyses were focused on *F_beta_*.

### Statistical analysis

Analyses were done in RStudio (RStudioTeam 2015) and the package *lme4* (Bates *et al*. 2015) and in JMP® 15.2.1. Pearson correlation coefficients (r) were used to test for correlations when visual inspection of the plots suggested that the conditions for parametric statistics were not significantly violated, else Spearman rank order correlations (rs) were used. Generalized linear models (GLM) and linear mixed models (LMM) were run on categorical or continuous dependent variables, respectively. GLM were based on a binomial error distribution. Models were reduced in stepwise regression with backward term selection in the built-in function *drop1*, and model comparison was done with likelihood-ratio test (LRT). The non-violation of the model assumptions was checked visually using fitted values versus residuals plot and residuals QQ-plots.

When testing for effects of ovarian fluids on sperm characteristics and fertilization success in the sperm competition trials, directed tests were used based on the *a priori* predictions that if ovarian fluids have an effect on sperm, the are expected to support sperm of more colorful and/or less inbred and/or less related males (the corresponding p-values are notated as p_dir_). Directed tests were used because such tests avoid inflation of the alpha value while still allowing to discover effects in the non-expected direction if the effect sizes are strong (Rice & Gaines 1994).

## Results

### Male phenotypes, leukocyte counts, MLH, and milt characteristics

Males looked silvery with variable intensity of ventral coloration. This coloration was striking on some individuals while largely absent on others. Colorful males were mostly yellow (Fig. 1), in few cases also slightly reddish (but 19 of 34 males had negative *a\** values, and the mean (±95% CI) *a\** value for the males was 0.95±1.78). Redness was positively correlated with yellowness (r = 0.45, p < 0.005) and therefore ignored in further analyses.

**Figure 1.**
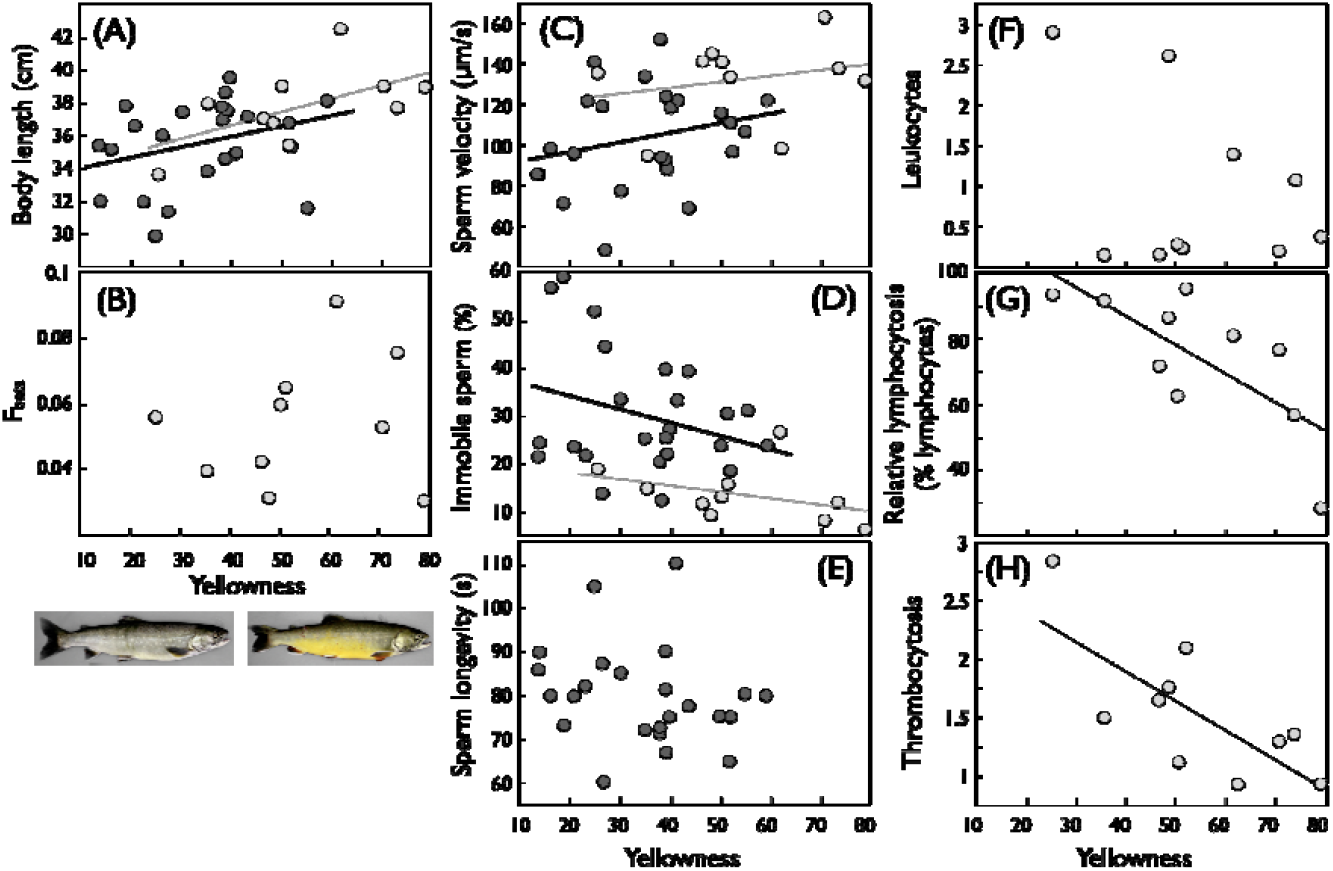
Yellowness in male lake char and their links to size, inbreeding coefficient, sperm characteristics, and immune parameters. The two fish illustrate the two extremes of observed yellowness in 2017. Yellowness versus (A) body size (r = −0.51, n = 34, p = 0.002), (B) inbreeding coefficients (F_beta_; r = 0.16, p = 0.65), the sperm characteristics (C) velocity, (D) rate of immobile sperm, and (E) longevity (see Table S3 for statistics on sperm characteristics), and the immune parameters (F) leukocyte counts (% of all blood cells; r = - 0.47, p = 0.17), (G) relative lymphocytosis (% of lymphocytes among leukocytes; r = −0.73, p = 0.02), and (H) thrombocytosis (thrombocytes counts per 100 blood cells; r = −0.74, p = 0.01). Regression lines illustrate significant relationships. Symbols and regressions in light grey indicate samples collected in 2017, darker symbols and regression lines those from 2018.

Yellowness increased with male size (Fig. 1A) but was not significantly correlated with *F_beta_* (Fig. 1B). Yellower males had less concentrated milt (LRT: *p* = 0.03; Table S4A). When their milt was activated in water only, yellower males had faster sperm (LRT: *p* = 0.05; Table S4B; Fig. 1C) and fewer immobile sperm than paler males (LRT: *p* = 0.05, Table S4C; Fig. 1D). Sperm longevity was not significantly correlated to skin coloration (LMM: p = 0.17, Table S4D; Fig. 1E). Including sampling year in our models explained 0% of the variance in sperm concentration but 5.1% and 15.8% of the variance in velocity and rate of immobile sperm, respectively. None of the sperm traits could be predicted by male size (Table S4) nor *F_beta_* except that males with higher *F_beta_* had higher counts of immobile sperm Fig. S4).

Some pale males had high leukocyte counts, but the correlation between leukocyte counts and skin coloration was not significant (Fig. 1F). However, skin coloration turned out to be a good predictor of relative lymphocytosis (Fig. 1G) and of thrombocytosis that both declined with increasing skin yellowness (Fig. 1H). No immune parameter was significantly correlated to male *F_beta_* (Fig. S4).

### Experiment 1: Sperm performance in ovarian fluid

The effects of activating sperm with water only or water with ovarian fluid can be analyzed on two levels, namely ignoring potential effects of male and female identity or including them. When ignoring male and female identities, activating sperm with or without ovarian fluids did not seem to affect average sperm velocity (LRT: χ^2^ = 1.01, *p* = 0.29, Fig. 2A). However, activating with ovarian fluids significantly reduced the variance in sperm velocity (Fig. 2A). Moreover, sperm activated with ovarian fluid were generally more motile (LRT: χ^2^ = 5.22, *p* = 0.022, Fig. 2B) and showed higher longevity (LRT: χ^2^ = 124.83, *p* < 0.001, Fig. 2C).

**Figure 2.**
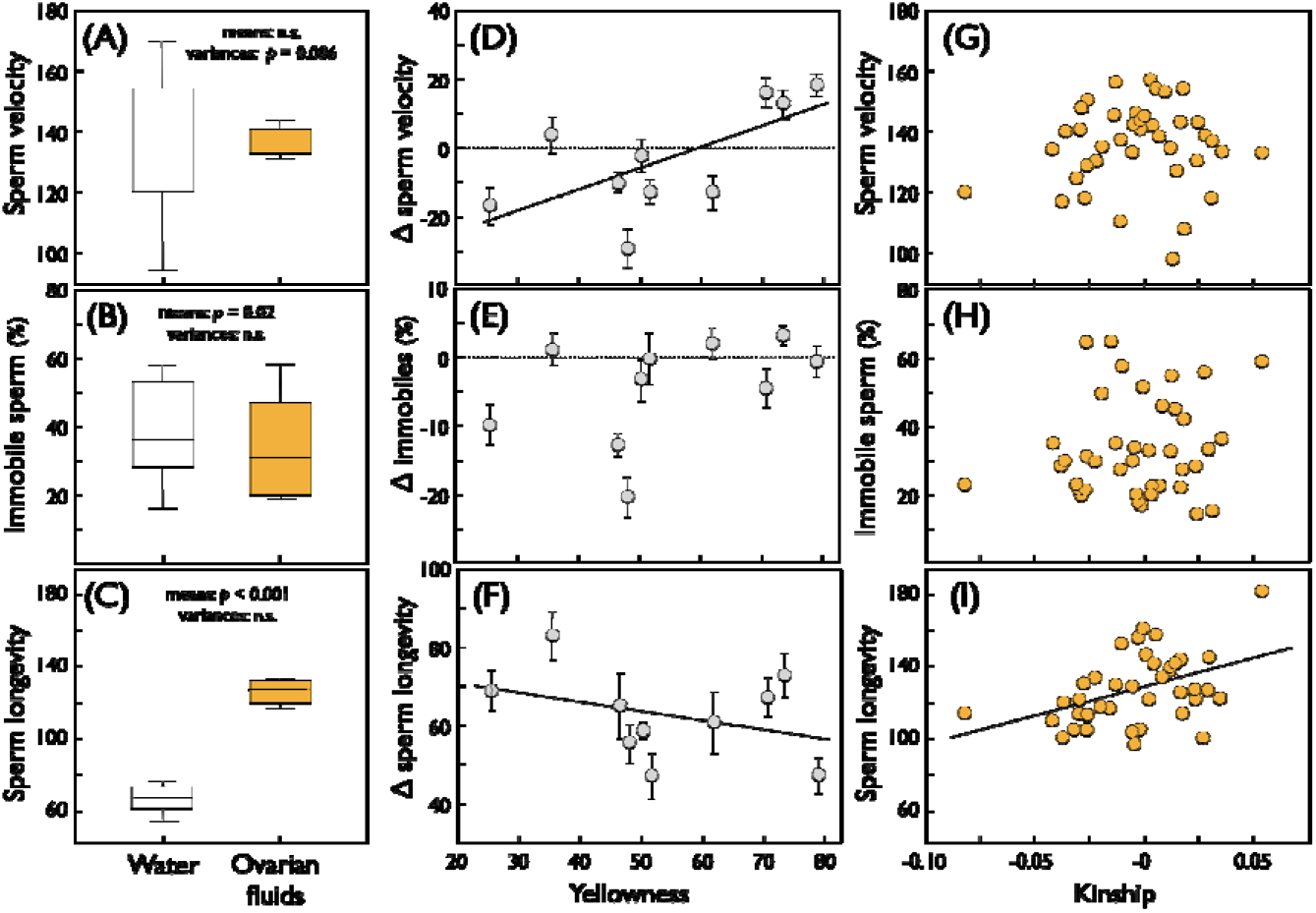
The effects of ovarian fluids on sperm of different types of males. Panels A-C show sperm characteristics (Turkey box plots with quartiles and whiskers) when activated in water only or in diluted ovarian fluids: (A) sperm velocity (μm/sec; comparing means: Welch’s F_1,12,014_ = 0.18, p = 0.68; variances: Levene’s F_1,18_ = 9.9, p = 0.006), (B) rate of immobile sperm (means: F_1,17.995_ = 0.48, p = 0.50; variances: F_1,18_ = 0.03, p = 0.87), and (C) sperm longevity (sec; means: F_1,14.486_ = 207.6, p < 0.001; variances: F_1,18_ = 0.5, p = 0.49). Panels e-g give the differences between of sperm characteristics in ovarian fluid and in water (8 measurements per male in ovarian fluids compared to the respective mean in water each) relative to male yellowness: (D) differences in sperm velocity (regression on means per male: r = 0.65, n = 10, p = 0.04), (E) differences in rates of immobile sperm (r = −0.36, p = 0.31), (F) differences in longevity (r = 0.65, p = 0.04). Panels G-I give the mean sperm characteristics per combination of male identity and donor of ovarian fluid relative to their kinship: (G) sperm velocity (r = 0.08, n = 40, p = 0.64), (H) rate of immobile sperm (r = 0.16, p = 0.34), and (I) sperm longevity (Spearman rank order correlation coefficient to account for potential extremes: rs = 0.39, p_two-tailed_ = 0.012, p_dir_ = 0.03 based on the *a priori* prediction that kinship should be avoided). Regression lines illustrate significant relationships, the dotted line in (D) and (E) mark zero difference.

When including potential effects of male and female identity, they both turned out to play significant roles. Males significantly varied in all sperm traits that were monitored here (see main effects of male identity in Table 1). Females significantly varied in how their ovarian fluids affected average sperm velocity and longevity but not the percentage of immobile sperm (see main effects of female identity in Table 1). Similar patterns could be observed with regard to how the sperm of different males responded to the ovarian fluids of different females (the male × female interaction terms in Table 1; Fig. S6). Again, average sperm velocity and longevity but not the percentage of immobile sperm depended on the combination of male and female identities (Table 1).

**Table 1.** Sperm characteristics after activation in water with different types of ovarian fluid (likelihood ratio tests on linear mixed-effects models). Significant *P*-values are highlighted in bold.

| Model terms | Effect tested | d.f. | AIC | $\chi^2$ | *p* |
| --- | --- | --- | --- | --- | --- |
| *(A) Sperm velocity* |  |  |  |  |  |
| h + m + f |  | 5 | 649.0 |  |  |
| m + f | h | 4 | 653.3 | 6.3 | **0.01** |
| h + f | m | 4 | 660.1 | 13.2 | **<0.001** |
| h + m | f | 4 | 652.6 | 5.7 | **0.02** |
| h + m × f | m × f | 6 | 637.8 | 13.2 | **<0.001** |
| *(B) Sperm motility* |  |  |  |  |  |
| m + f |  | 4 | 587.6 |  |  |
| f | m | 3 | 670.1 | 84.6 | **<0.001** |
| m | f | 3 | 585.6 | 0 | 1 |
| m × f | m × f | 5 | 589.6 | 0 | 1 |
| *(C) Longevity* |  |  |  |  |  |
| m + f |  | 4 | 664.7 |  |  |
| f | m | 3 | 687.7 | 25.0 | **<0.001** |
| m | f | 3 | 694.2 | 31.5 | **<0.001** |
| m × f | m × f | 5 | 527.6 | 139.1 | **<0.001** |

Fixed effect: h, sperm head size; random effects: m, male identity; f, female identity

Sperm responses to ovarian fluids could partly be predicted by male skin coloration. With increasing yellowness, sperm velocity increased more in the presence of ovarian fluids (Fig. 2D). The reduction of immobile sperm that ovarian fluids caused could not be predicted by male yellowness (Fig. 2E), but the increase in sperm longevity in response to ovarian fluids was less pronounced in more yellow males than in pale ones (Fig. 2F).

The 40 different combinations of male identity and donor or ovarian fluid allowed to test whether the genomic distance between males and female (kinship) affected sperm performance in ovarian fluid. Kinship seemed to have no effects on sperm velocity or percentage of immobile sperm (Fig. 2G,H). However, kinship between male and female increased the longevity of sperm when exposed to ovarian fluid (Fig. 2I).

### Experiment 2: Males success in sperm competition

We genotyped 1,475 offspring of which 1,464 could be assigned to their father (99.3% success). In the following, three types of focal males are considered: the most yellow, the least inbred, and the least related to the respective female, because male yellowness did not correlate with their inbreeding coefficients (Fig. 1B; p = 0.65) nor with kinship (Fig. S4A; p = 0.42), and kinship could not be predicted by male inbreeding coefficients (Fig. S4B; p = 0.47).

Figure 3 shows the competitive success of different types of focal males when tested in water only or with ovarian fluids (including only the females for which we had both types of conditions; see Table S5 for the identities of the focal males in the different scenarios). Overall, the success rates of focal males were 53.6%, 56.2%, and 53.2% if the focal male was the most yellow, the least inbred, or the least related to the female, respectively (Fig. 3). Dyad identity always had a strong effect on the focal male’s success rate (Table 2). The presence of ovarian fluid significantly enhanced the focal male’s success if it is the yellower of the two competitors (Table 2A, Fig. 3A) while female identity or the difference in sperm velocity between the rivals played no significant roles in this scenario (Table 2A). Similar patterns could be observed if the focal male was the one with the lower F_beta_, except that the presence of ovarian fluid did not increase the focal male’s fertilization success (Table 2B, Fig. 3B). No effects of ovarian fluids could be observed if the focal male was the least related to the egg donor (Table 2C, Fig. 3C). In the latter scenario, the female identity and the difference in sperm velocity now explained male success (Table 2C).

**Figure 3.**
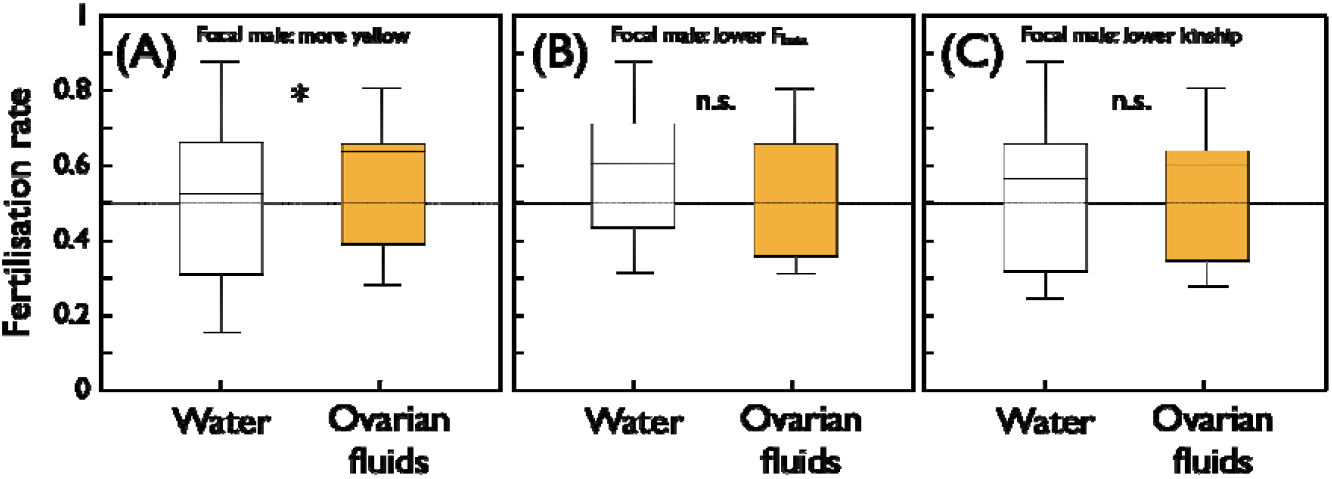
Competitive success of different types of focal males under sperm competition in water only or when exposed to ovarian fluids. The mean fertilization rates of (A) the yellower male of the two competitors each, (B) the less inbred male each, and (C) the male each that is least related to the female whose ovarian fluids the sperm was exposed to. Turkey box plots with quartiles and whiskers (N_water_ = N_ovarian fluid_ = 15). The asterisk indicates a significant difference linked to the activation medium, n.s. = not significant. See Table 2 for statistics.

**Table 2.** Correlates to fertilization success of the focal male in the sperm competition trials (experiment 2), if the focal male is (A) the more yellow one of the two competitors, (B) the one with the lower inbreeding coefficient *F_beta_*, or (C) the least related to the female. Generalized linear model with female (egg donor), dyad (pair of competing males), ovarian fluid (present of absent), and the difference in mean sperm velocity between rivals (focus – opponent, in water or in ovarian fluid, respectively, as determined in experiment 1). Nonsignificant interaction terms were removed. Final tests were either two-tailed or directed (*p_dir_*) based on the *a priori* predictions that ovarian fluids have either no effect or they support sperm of focal males. Final *P*-values are highlighted in bold if significant.

| Factor | d.f. | $\chi^2$ | $p_{two-tailed}$ | $p_{dir}$ |
| --- | --- | --- | --- | --- |
| *(A) Focal male: more yellow* |  |  |  |  |
| Female | 2 | 3.1 | 0.21 |  |
| Dyad | 4 | 130.7 | **<0.001** |  |
| Ovarian fluid | 1 | 4.3 | 0.039 | **0.02** |
| Difference in sperm velocity | 1 | 0.8 | 0.37 | 0.23 |
| *(B) Focal male: lower $F_{beta}$* |  |  |  |  |
| Female | 2 | 1.0 | 0.60 |  |
| Dyad | 4 | 119.2 | **<0.001** |  |
| Ovarian fluid | 1 | 4.2 | 0.040 | 0.10 |
| Difference in sperm velocity | 1 | 0.9 | 0.35 | 0.22 |
| *(C) Focal male: lower kinship* |  |  |  |  |
| Female | 2 | 27.0 | **<0.001** |  |
| Dyad | 4 | 57.8 | **<0.001** |  |
| Ovarian fluid | 1 | 0.1 | 0.76 | 0.48 |
| Difference in sperm velocity | 1 | 8.8 | 0.003 | **0.002** |

## Discussion

The first question of this study was whether sperm traits of wild-caught lake char can be predicted from male phenotypes (body size and skin coloration) or overall genetic quality (here determined as F_beta_) when tested in water only, independently of any potential effects of ovarian fluids. We found no effects of body size, but skin coloration was a good predictor of sperm traits: yellower males had faster sperm and lower rates of immobile sperm (i.e. higher sperm motility) than paler males. The skin coloration of lake char is hence among the secondary sexual traits that have been found to be positively linked to sperm competitiveness in different fish families (e.g. Locatello *et al*. 2006; Janhunen *et al*. 2009; Mehlis *et al*. 2013; Fukuda & Karino 2014). Our findings also suggest that paleness in our sample was linked to comparatively low health and vigor rather than to a “sneaker” life-history strategy (parasitic male spawners) that are common in many migratory salmonids, because sneakers are predicted to invest more into sperm than dominant males (Ball & Parker 2000).

Secondary sexual traits like conspicuous skin colors are predicted to reveal general health and vigor (Andersson 1994). This has repeatedly been observed in other species (Milinski & Bakker 1990; Johnson & Fuller 2015). In our study, paler males suffered from increased relative lymphocytosis and thrombocytosis, i.e. they had increased rates of lymphocytes and of thrombocytes, two potential indicators of acute infections or other physiological stress (Haenen *et al*. 2010; Rose *et al*. 2012; Yan *et al*. 2013), than yellower males. Male yellowness was hence a useful predictor of current male health condition, potentially revealing, for example, virulence of a tapeworm infection (Johansen *et al*. 2019).

Theory also predicts a link between F_beta_ and general health and vigor (Allendorf & Luikard 2007), and such links have indeed been found in other studies (Fox & Reed 2011). We found that the rates of immobile sperm declined with declining male F_beta_, supporting the hypothesis that sperm motility can be an indicator of general health and vigor (Kowalski & Cejko 2019). We did not find any significant correlations between male F_beta_ and immune parameters, but this may need to be studied again in larger samples that provide more statistical power.

Despite the similar expectations from theory, F_beta_ and skin colorations turned out to be not significantly correlated in our sample. In previous studies, such a correlation was sometimes found (e.g. Zajitschek & Brooks 2010; Herdegen *et al*. 2014) and sometimes not (e.g. Frommen *et al*. 2008; Marsh *et al*. 2017), including a study on Arctic char (Janhunen *et al*. 2011). One possibility is that skin coloration is a plastic trait that quickly reacts to acute stress (Johnson & Fuller 2015) while F_beta_ affects more the overall tolerance to environmental stress. If so, the link between F_beta_ and skin colorations is expected to change with changing environmental conditions.

The next major questions of this study were whether and how ovarian fluids affect sperm traits, and whether such effect differ for different types of males. We found that ovarian fluids affected all sperm traits that we studied: They reduced the between-male variance in sperm velocity, reduced the percentage of immobile sperm, and nearly doubled mean sperm longevity. Sperm velocity was not generally increased by ovarian fluids, apparently contrary to some previous findings in other fishes (Zadmajid *et al*. 2019). However, when comparing the effects of ovarian fluids on sperm characteristics of different types of males, it turned out that the yellower a male, the more does the velocity of its sperm increase when in contact with ovarian fluids, while sperm of paler males are rather slowed down in ovarian fluids. Such a correlation between the effects of ovarian fluids and male coloration suggests that sperm of yellower males are better prepared to ovarian fluids while sperm of paler males are better prepared to the absence of ovarian fluids. If yellower males are closer to the female vent during spawning than paler males, these reaction of sperm to the different environments correspond well to the expected microenvironment that the sperm are likely to be exposed to.

The effects of ovarian fluids on sperm motility was not correlated to male yellowness, i.e. all male types profited similarity from the presence of ovarian fluids. Interestingly, however, sperm longevity was more enhanced by ovarian fluids in paler males than in yellower males, suggesting that there is a trade-off between increased velocity and increased longevity. Sperm of yellower males profited from increased velocity while sperm of paler males profited from increased longevity when in contact with ovarian fluids.

Females varied in how much their ovarian fluids generally increased sperm velocity and sperm longevity, but not in how much their ovarian fluids reduced sperm motility. Females also varied in how much their ovarian fluids specifically increased velocity and longevity of sperm of certain males over others, i.e. we found the corresponding females x male interactions to be significant (but again not in the case of sperm motility). When we tested whether some of these interactions can be explained by the genetic distance between males and females, we found no link between relatedness and sperm velocity or motility, and a non-expected positive link between relatedness and sperm longevity. Ovarian fluids seemed to specifically elongate the longevity of sperm of more closely related males. This non-expected finding needs to be further studied. We can currently not offer an adaptive explanation for why females should support the longevity of sperm of related males more than that of sperm of unrelated males. Preferring unrelated males would reduce the average offspring inbreeding coefficients.

Our next major question was: Whose sperm benefit from exposure or non-exposure to ovarian fluids when in sperm competition? Here we focused on three types of males: the yellower ones of the experimental dyads, the less inbred ones, and the ones who were least related to a given female. We found that the presence of ovarian fluids led to enhanced fertilization success of the more yellow male but did not specifically affect the sperm of less inbred or less related males. However, competition trials can only reveal a limited range of possible situations. In our case, the overall sperm to egg ratios were so high that variation in sperm longevity may not have affected the outcome of the sperm competition, while late fertilizations may often be relevant under natural conditions.

In a parallel study, Garaud et al. (2022) used an extended sample of our study population to test whether females would profit from mating with more yellow, less inbred, or less related males. They used a full-factorial breeding design (crossing 10 male and 6 females in all possible combinations) and single rearing of large numbers of offspring to test the embryos’ reaction to the pathogen *Aeromonas salmonicida*, to an ecologically relevant concentration of a common chemical pollutant (a spike of 1 ng/L ethinylestradiol into the water), and to no-stress conditions (sham controls). They found that while exposure to ethinylestradiol showed no effects, exposure to the pathogen reduced embryo development rate. Contrary to expectations from ‘good-genes’ hypotheses of sexual selection, this pathogen-related reduction in embryo development rate increased with increasing male yellowness, i.e. offspring of yellower males were on average smaller and less tolerant to the pathogen stress than offspring of paler males. These results confirm previous findings on Arctic char (Janhunen *et al*. 2011) and suggest that females would not profit from giving yellower males a selective advantage during sperm competition. However, Garaud et al. (2022) found strong effects of the relatedness between males and females on offspring performance. Embryo development rates declined with increasing parental relatedness and hence increasing mean inbreeding coefficients among the offspring. Similar effects of relatedness could be observed in the embryos and larvae that resulted from experiment 2 of the present study and that are also presented in detail in Garaud et al. (2022), while male yellowness or male F_beta_ were both not significantly linked to the growth of these embryos (Garaud *et al*. 2022).

In conclusion, male yellowness revealed aspects of current health and vigor and was generally linked to high sperm velocity and motility. If exposed to diluted ovarian fluids, sperm velocity of yellower males became even faster and sperm of paler males slowed further down. In sperm competition experiments, fertilization success of the yellower of the two competitors was indeed increased when exposed to diluted ovarian fluids. Exposure to ovarian fluids also generally increased sperm motility and longevity. While motility was similarity increased in sperm differently colored males, longevity was more enhanced in sperm of paler males than in sperm of yellower males, suggesting a trade-off between increased sperm velocity and increased sperm longevity. Male F_beta_ and male genetic relatedness to a given female were poor predictors of sperm performance in water or ovarian fluids. When offspring of the different types of males were tested under various types of stress conditions, offspring of yellower males grew slower and were less stress tolerant than offspring of paler males, while offspring of least related parents showed the best performance in the various stress conditions. Combined, these results suggest that the increased performance of sperm of yellower males in ovarian fluids is an adaptation to the microenvironment that the sperm are likely to be exposed to, rather than cryptic female choice by ovarian fluids.

## Ethics

This work complied with the national, cantonal and university regulations where it was carried out. The handling and transport of adults and transport of embryos was approved by the French authorities (INTRA.FR.2017.0109258).

## Data accessibility

Data will be deposited on the Dryad depository upon acceptance of the manuscript.

## Acknowledgments

We thank G. Levray from the Pistolet fishery for catching the fish, the APERA association and the *pisciculture des Rives* for access to the fish, L. Adhia Eya, A. Atherton, C. Berney, J. Bideaux, F. Dolivo, L. Espinat, S. Kreuter, E. Longange, L. Marques da Cunha, and N. Sironi for assistance in the field or in the lab, F. Schütz for discussion on statistics, and E. Lasne for discussion and access to the INRA facilities.

## Supplementary information

**Table S1.**
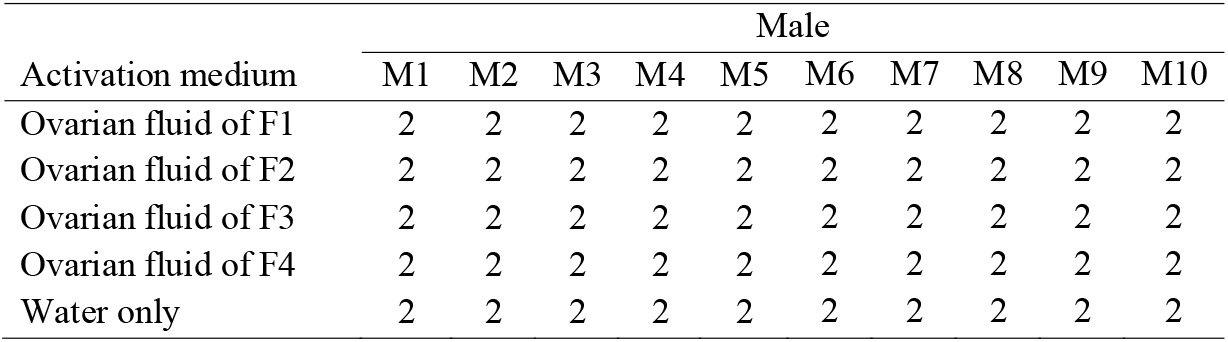
Experiment 1: Determining sperm characteristics in various activation media, i.e. diluted ovarian fluid of one of 4 females (F1-F4) or water only. The entries give the number of independent sets of measurements.

**Table S2.** Experiment 2: The sperm competition trials, with 2 times 24 eggs per experimental cell. Asterisks indicate experimental cells that were accidentally lost.

| Female | Activation | Dyad |  |  |  |  |
| --- | --- | --- | --- | --- | --- | --- |
|  |  | M1 vs M2 | M3 vs M4 | M5 vs M6 | M7 vs M8 | M9 vs M10 |
| F1 | ovarian fluid | 2 x 24 | 2 x 24 | 2 x 24 | 2 x 24 | 2 x 24 |
| F1 | water | 2 x 24 | 2 x 24 | 2 x 24 | 2 x 24 | 2 x 24 |
| F2 | ovarian fluid | 2 x 24 | 2 x 24 | 2 x 24 | 2 x 24 | 2 x 24 |
| F2 | water | 2 x 24* | 2 x 24* | 2 x 24* | 2 x 24* | 2 x 24* |
| F3 | ovarian fluid | 2 x 24 | 2 x 24 | 2 x 24 | 2 x 24 | 2 x 24 |
| F3 | water | 2 x 24 | 2 x 24 | 2 x 24 | 2 x 24 | 2 x 24 |
| F4 | ovarian fluid | 2 x 24 | 2 x 24 | 2 x 24 | 2 x 24 | 2 x 24 |
| F4 | water | 2 x 24 | 2 x 24 | 2 x 24 | 2 x 24 | 2 x 24 |

**Table S3.**
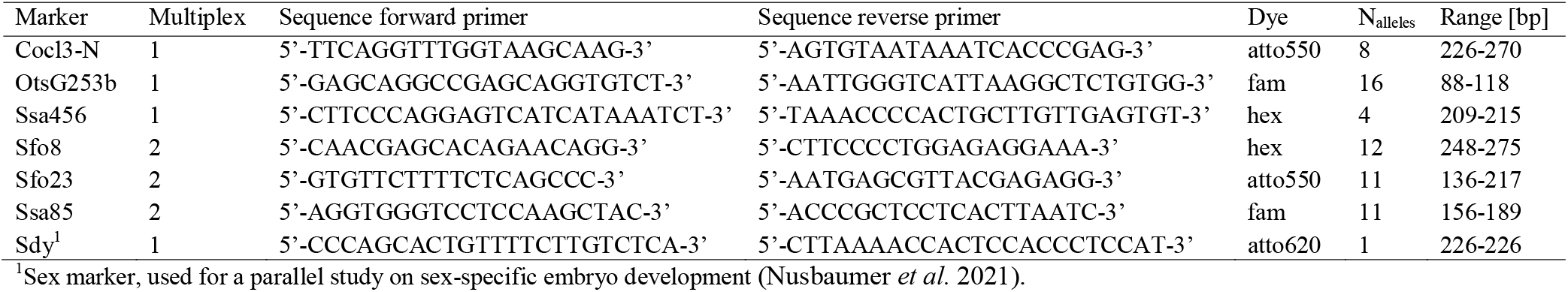
The 6 microsatellite markers used for paternity assignment, and the sex marker that was added to the first multiplex.

**Table S4.**
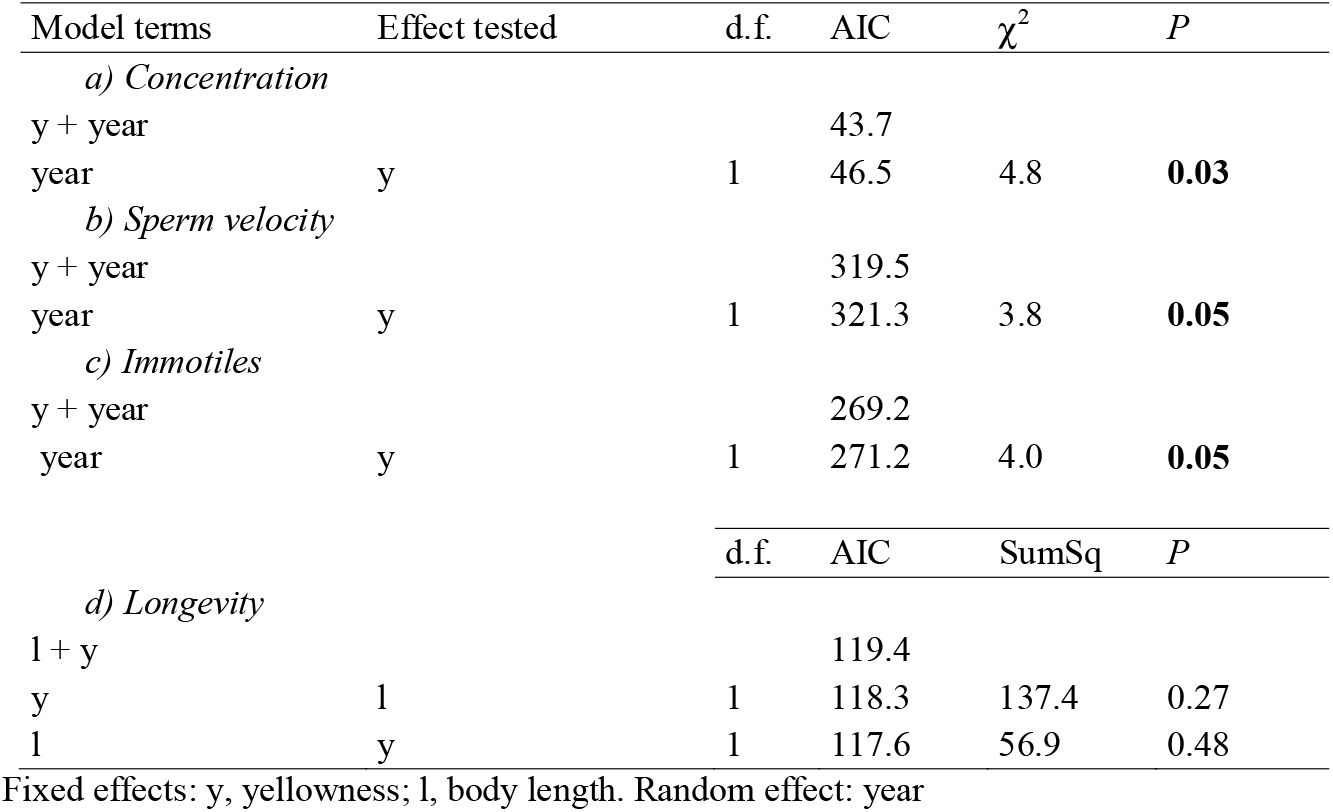
Characteristics of sperm activated in water only, predicted by male body length, yellowness and year of sampling. Likelihood-ratio tests on linear mixed-effects models were used when year of sampling could be added as random factor (a-c), while linear models were used when measurements were only taken in 2017 (d). The table shows the outcome after removal of least significant parameter and gives the AIC for each model. Bold *P*-values highlight significant effects (*P* ≤ 0.05).

**Table S5.** Identity of focal males if it is the more yellow than its competitor, less inbred, or least kin to a given female.

| Dyad | More yellow | Less inbred | Least kin to female |  |  |  |
| --- | --- | --- | --- | --- | --- | --- |
|  |  |  | F1 | F2 | F3 | F4 |
| 1 | M2 | M2 | M2 | M2 | M2 | M1 |
| 2 | M3 | M4 | M4 | M3 | M4 | M3 |
| 3 | M5 | M6 | M5 | M6 | M6 | M6 |
| 4 | M7 | M8 | M8 | M8 | M8 | M8 |
| 5 | M10 | M9 | M10 | M9 | M10 | M10 |

**Figure S1.**
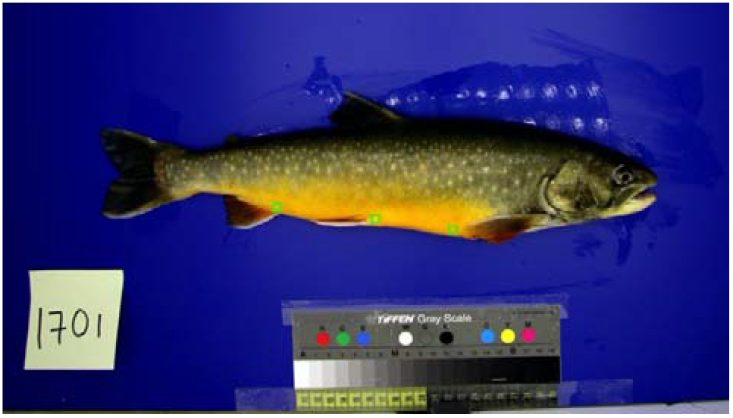
Locations of the 3 squares (in green) from which color measurements of the ventral area (yellow and red) were taken. Mean gray value was positively correlated with yellowness, i.e. the yellower the lighter (r = −0.47, p < 0.005) and negatively with redness, i.e. the redder the darker (r = −0.66, p < 0.001).

**Figure S2.**
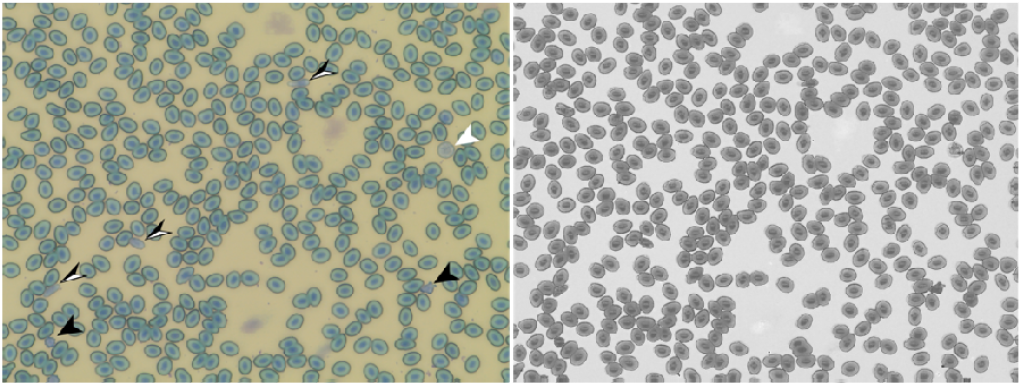
Example of a blood smear image as taken under the microscope (left) and its output of automated cell count (right). On the left image, black arrow heads point towards lymphocytes, the white arrow towards a granulocyte, and the black and white arrow towards thrombocytes. The rest are erythrocytes.

**Figure S3.**
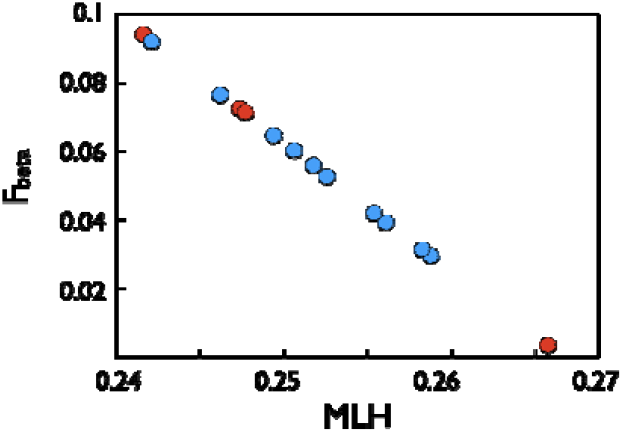
The link between multi-locus heterozygosity (MLH) and individual inbreeding coefficients (*F_beta_*) for the males (blue) and females (red) sampled in 2017 and used in experiments 1 and 2.

**Figure S4.**
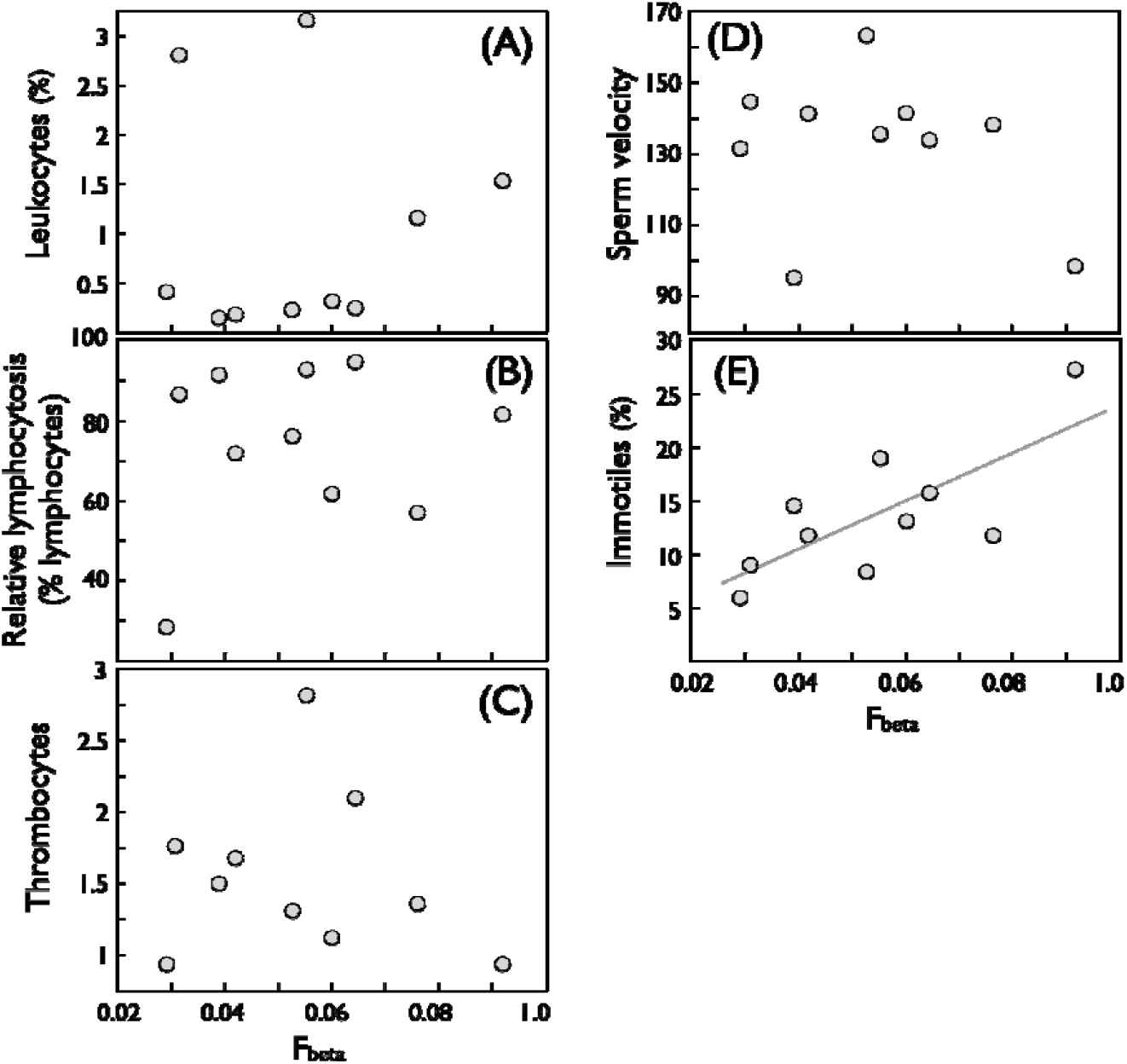
Immunological and sperm characteristics of males relative to their inbreeding coefficient (F_beta_): (A) leukocytes (% of all blood cells; r = 0.17, p = 0.88), (B) relative lymphocytosis (percentage of lymphocytes among leukocytes; r = 0.11, p = 0.61), (C) thrombocytes per 100 blood cells (r = −0.20, p = 0.75), (D) sperm velocity (μm/sec; r = −0.25, p = 0.48), (E) percentage of immotile sperm (r = 0.74, p = 0.01). The regression line is given for the significant relationship. Sperm longevity was not measured in this sample that was taken in 2017.

**Figure S5.**
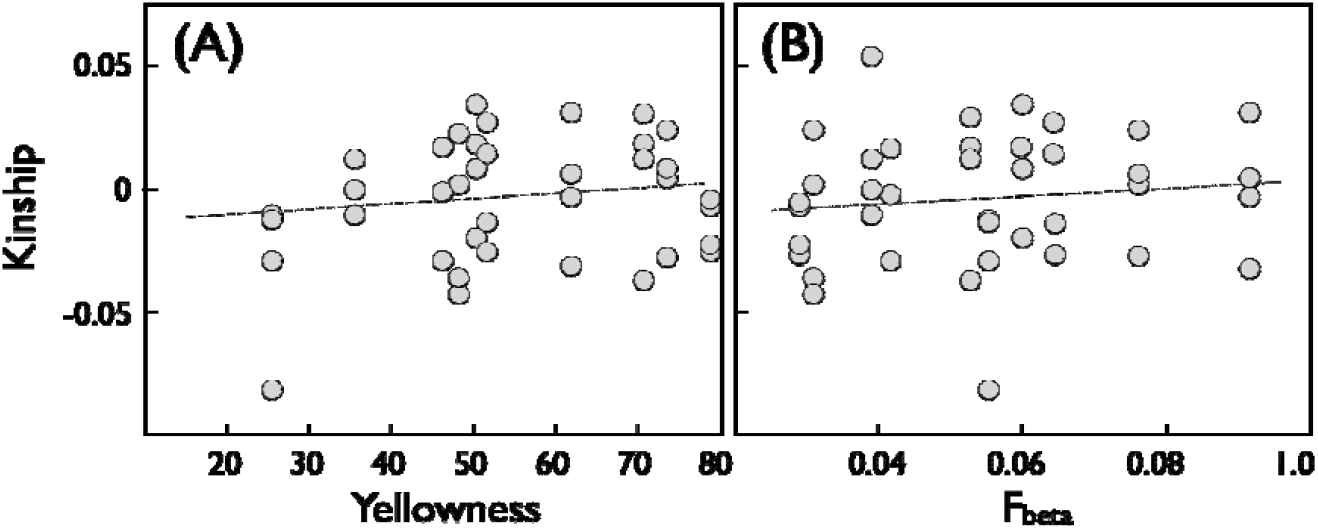
Relatedness between males and females (kinship) that were used in the sperm competition trials (4 females x 10 males = 40 combinations) versus (A) male skin colorations (yellowness; r = 0.13, n = 40, p = 0.42) and (B) individual male inbreeding coefficients (*F_beta_*; r = 0.12, p = 0.47). The dotted lines indicate the non-significant regressions.

**Figure S6.**
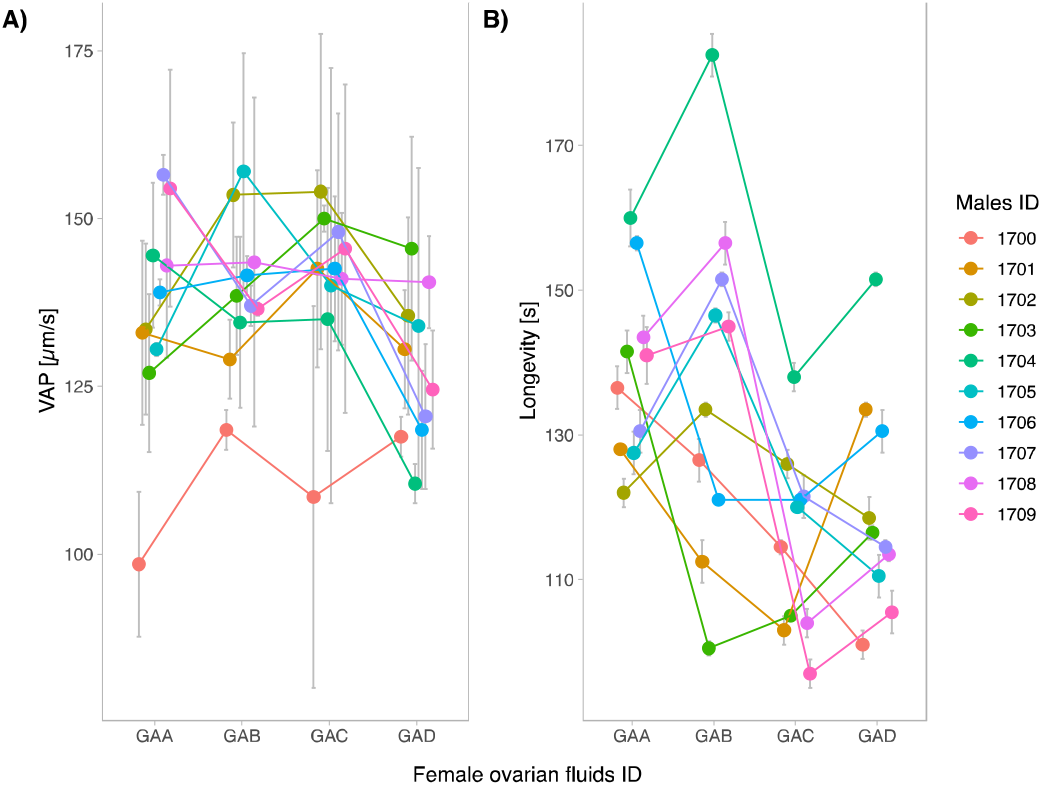
Interactions effects between females’ ovarian fluids and males’ identities on A) sperm velocity and (B) sperm longevity. Symbols depicts means of within subject repeated measures with their 95% confidence intervals. See Table 1 for statistics.

